# Chronic environmentally relevant levels of Simvastatin induces non-monotonic responses in Zebrafish (*Danio rerio*)

**DOI:** 10.1101/289694

**Authors:** Susana Barros, Rosa Montes, José Benito Quintana, Rosario Rodil, Ana André, Ana Capitão, Joana Soares, Miguel M. Santos, Teresa Neuparth

**Affiliations:** CIMAR/CIIMAR – Interdisciplinary Centre of Marine and Environmental Research, Endocrine Disruptors and Emerging Contaminants Group, University of Porto, Avenida General Norton de Matos, S/N 4450-208 Matosinhos, Portugal; Department of Analytical Chemistry, Nutrition and Food Sciences, IIAA — Institute for Food Analysis and Research, University of Santiago de Compostela, Constantino Candeira S/N, 15782 Santiago de Compostela, Spain; FCUP – Department of Biology, Faculty of Sciences, University of Porto (U.Porto), Porto, Portugal

**Keywords:** Zebrafish, Simvastatin, *hmgcr*, Embryonic development, Cholesterol and Triglycerides, Chronic effects, Low-level exposures

## Abstract

Simvastatin (SIM), a hypocholesterolaemic compound, is among the most prescribed pharmaceuticals for cardiovascular disease prevention worldwide. Several studies have shown that acute exposure to SIM is able to produce multiple adverse effects in aquatic organisms. However, uncertainties still remain regarding the chronic effects of SIM in aquatic ecosystems. Therefore, the present study aimed to investigate the effects of SIM in the model freshwater teleost zebrafish (*Danio rerio*) following a chronic exposure (90 days) to environmentally relevant concentrations ranging from 8 ng/L to 1000 ng/L. This study used a multi-parametric approach integrating distinct ecological-relevant endpoints, i.e. survival, growth, reproduction and embryonic development, with biochemical markers (cholesterol and triglycerides). Furthermore, Real Time PCR was used to analyse the transcription levels of key genes involved in the mevalonate pathway (*hmgcra, cyp51*, and *dhcr7*). Globally, SIM induced several non-monotonic dose-responses; embryonic development, biochemical and molecular markers, were significantly impacted in the low-intermediate concentrations, 40 ng/L and 200 ng/L, whereas no effects were recorded for the highest tested SIM levels (1000 ng/L). Taken together, these findings expand our understanding of statins effects in teleost’s, demonstrating significant impacts at environmentally relevant concentrations. The findings highlight the importance of addressing the effects of chemicals under chronic low-level concentrations.

**Graphical abstract:** 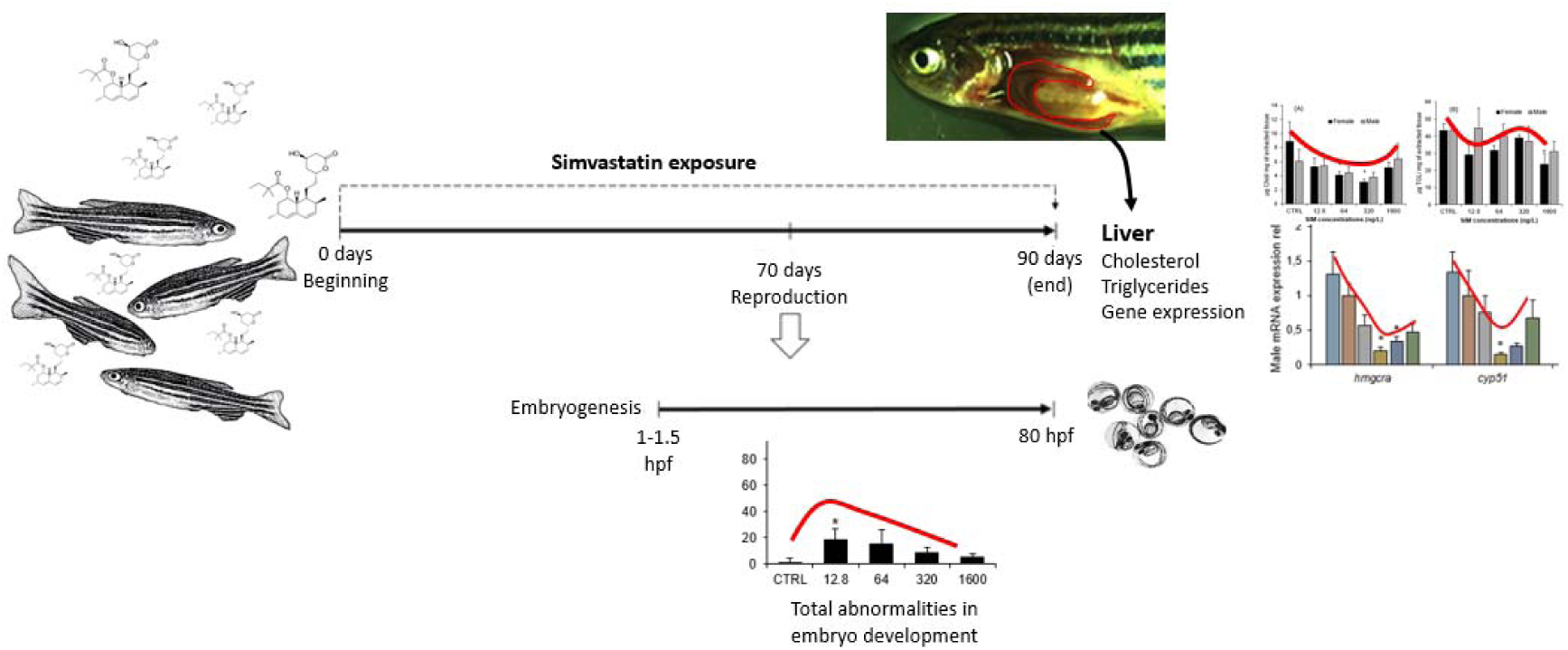

**Highlights:**

- Several uncertainties exist regarding simvastatin mode of action in non-target organisms
- This work integrates *D. rerio* multi-level responses after long-term exposure to simvastatin
- Simvastatin impacted cholesterol/triglycerides levels and transcript levels of genes related to mevalonate pathway.
- Parental exposure to simvastatin induced offspring embryonic malformations.
- Embryonic abnormalities, biochemical and molecular data did follow a non-monotonic curve.

## 1. Introduction

In the past, pharmaceuticals were overlooked as aquatic pollutants because exposure levels were considered to be too low to induce significant effects in non-target organisms (Arnold et al., 2014; Daughton, 2016; EEA, 1999). However, from the mid-90s, a growing attention has been devoted to this class of compounds. Indeed, the detection of pharmaceuticals in the aquatic environments has increased in the last years, not only because of the pharmaceutical industry growth, but also due to improvements of analytical methods (LaLone, et al., 2014). Most pharmaceuticals are detected in surface waters at trace levels, generally, at concentrations ranging between ng/L to the low μg/L levels (Arnold et al., 2014; Azzouz & Ballesteros, 2012; BIO Intelligence Service, 2013; Daughton, 2016; Fent et al., 2006). However, since they are bioactive substances, designed to produce biological effects at rather low concentrations, the scientific community is now in broad agreement that pharmaceuticals may pose a considerable environmental risk (Rodrigues et al. 2006, Ferreira et al., 2009; BIO Intelligence Service, 2013). In fact, several pharmaceuticals have been demonstrated to induce effects at environmentally relevant concentrations in non-target organisms (Arnold et al., 2014; Fent et al., 2006; Neuparth et al., 2014). Nevertheless, the ecological risk assessment of pharmaceuticals is still on its infancy, because most studies dealing with this class of compounds only report acute toxicity effects, while these compounds concentrations in the wild are too low to pose acute toxicity risk (Dahl et al., 2006, Fent et al., 2006, Sárria et al., 2011). Hence, there is a need to assess chronic effects of pharmaceuticals given that non-target aquatic organisms are continuously exposed to low levels of this class of compounds for several generations (Arnold et al., 2014; BIO Intelligence Service, 2013; Fent et al., 2006).

Simvastatin (SIM) is a hypolipidemic drug of the statins class used in humans as the primary treatment of hypercholesterolemia to decrease serum LDL-cholesterol levels (Burg & Espenshade, 2011; Igel, et al., 2001; Neuparth et al., 2014; Vázquez et al., 2017). SIM, as well as other statins, is known to specifically inhibit the enzyme 3-hydroxy-3-methyl-glutaryl-CoA reductase (HMGCR), which is essential for *de novo* synthesis of cholesterol in mammalian’ cells through the mevalonate pathway (Al-Habsi, et al., 2016; Fent, et al., 2006; Blumenthal, 2000; Endo et al., 1976a; Endo, et al., 1976b; Sehayek, et al., 1994). SIM compete with HMG-CoA for the active binding site in the enzyme HMGCR, having an affinity of about three orders of magnitude greater than the natural substrate. Once bound to the enzyme, statins alter its conformation, inhibiting its function and thereby decreasing the cholesterol synthesis (Al-Habsi et al., 2016; Istvan, 2003; Moghadasian, 1999).

SIM has been reported to be among the most prescribed human pharmaceuticals in the western countries. Due to the lack of mechanisms that ensure complete removal of statins in wastewater treatment plants (WWTP), significant amounts of this pharmaceutical are discharged in the aquatic environments (Fent et al., 2006; Lapworth et al., 2012). Several authors have reported the presence of SIM in WWTPs worldwide (Kasprzyk-Hordern et al., 2009; Miao & Metcalfe, 2003; Ottmar et al., 2012; Pereira et al., 2016; Pereira et al., 2015; Sousa, 2013; Verlicchi et al., 2012). Concentrations up to 8.9 µg/L and 1.23 µg/L (influents) and 1.5 µg/L and 90 ng/L (effluents), were reported in Portugal and USA WWTPs, respectively (Ottmar et al., 2012; Pereira et al., 2016). The predicted environmental concentration for SIM in Norwegian and Portuguese surface waters have been estimated at 630 ng/L and 369.8 ng/L, respectively (Grung et al., 2007; Pereira et al. 2015).

Considering that SIM is being extensively used in the treatment of hypercholesterolemia, reaching the aquatic environment in increasing concentrations, many aquatic taxa might be at risk. This is particularly true given that statins were predicted to inhibit HMGR in a board range of animal *taxa* (Santos et al., 2016). In fact, the main concern of the presence of SIM in the aquatic ecosystems is its environmental persistence, toxicity, and bioactivity, at rather very low concentrations. Furthermore, SIM has a high log K_ow_ of 4.68, which might indicate high bioaccumulation potential in aquatic organisms (Santos, et al., 2016). Previous studies have reported several detrimental effects of SIM in aquatic organisms at several levels of biologic organization, such as impairment of embryo development (*Danio rerio* and *Parachentrotus lividus* – Ribeiro et al., 2015), decreased metabolic activity and membrane stability (*Oncorhynchus mykiss* hepatocyte – Ellesat, et al., 2010), alteration of acetylcholinesterase and lipid peroxidation levels (*Fundulus heteroclitus* – Key et al., 2009; *Palaemonetes pugio* – Key et al., 2008), disturbances in growth (*Dunaliella tertiolecta, Nitocra spinipes and Gammarus locusta* – Dahl et al., 2006; DeLorenzo & Fleming, 2008; Neuparth et al., 2014), severe impairment of reproduction (*Gammarus locusta* – Neuparth et al., 2014), and even mortality (*Plaemonetes pugio* – Key et al., 2008). With the exception of the Neuparth et al (2014) study, that reported chronic effects of SIM in the ng/L range, all the other studies mentioned above have been based on acute toxicity tests, with SIM concentrations above environmentally relevance.

Despite the diversity of studies regarding SIM toxicity, the underlying mechanisms of action in aquatic organisms are still not fully understood (Gee et al., 2015). Acknowledge the mode of action (MOA) of SIM in non-target aquatic organisms is important to build its ecotoxicological response and to anticipate the effects of compounds acting through a similar mechanism of action. This is particularly relevant considering that pharmaceuticals, such as SIM, are bioactive molecules designed to produce biological effects at very low concentrations. Hence, there is an urgent need to perform long-term, low level exposure assays, that integrates effects in ecologically-relevant endpoints with molecular and biochemical responses.

Therefore, the main aim of the present study was to evaluate SIM effects on zebrafish, following a chronic exposure to environmentally relevant concentrations (ng/L). We integrate multiple key ecological endpoints (survival, growth, reproduction, and embryonic development), with biochemical markers of lipid homeostasis (cholesterol and triglycerides) and molecular analysis (expression of key genes coding for the mevalonate pathway: *hmgcra, cyp51*, and *dhcr7*), to get insights into SIM long-term adverse effects in ecologically relevant endpoints and address the underlying mechanism of toxicity in fish.

## 2. Material and Methods

### 2.1 Species Selection

Zebrafish *(Danio rerio)* is recommended as a test species in a wide range of ecotoxicological test protocols (Oberemm, 2000). Its small size, robustness, multiple progeny from a single mating, embryos transparency and easy maintenance under laboratory conditions are advantages for its use as test organism (Fang & Miller, 2012; Lawrence, 2007; Soares et al., 2009). In addition, the close phylogeny of zebrafish and mammals, with highly conserved mevalonate pathway, makes this species ideal for performing the present study.

### 2.2. Zebrafish maintenance

Wild-type zebrafish, 50 days age-old, were obtained from Singapore local suppliers. Animals were acclimated to controlled laboratory conditions, in 250 L aquarium with dechlorinated filtered and aerated water. During this period, fish were kept at 28 ± 1°C, under a photoperiod of 14:10 h (light:dark) and fed, *ad libitum*, three times per day with commercial fish diet Tetramin (Tetra, Melle, Germany). These conditions were maintained for 15 days until the beginning of the chronic bioassay.

### 2.3. Chronic toxicity bioassay

The chronic bioassay was carried out at “Biotério de organismos aquáticos” (BOGA) located at Interdisciplinary Centre of Marine and Environmental Research (CIIMAR). The experiment was subject to a previous ethical review process carried out by CIIMAR animal welfare body (ORBEA). The bioassay was performed in compliance with the European Directive 2010/63/EU, on the protection of animals used for scientific purposes, and the Portuguese “Decreto Lei” 113/2013.

The bioassay lasted for 90 days, starting with the exposure of sub-adults zebrafish allocated in 30 L aquaria, under a flow-through system. The water flow was maintained at 1.08 L per hour by means of a peristaltic pump (ISM 444, ISMATEC) provided with dechlorinated heated and charcoal filtered tap water. Each aquarium, with 25 animals, was maintained with a water temperature of 28 ± 1°C, 14:10 h (light:dark) photoperiod and a mean ammonia concentration of 0.08 ± 0.04 mg/L. Fish were fed twice a day with a commercial fish diet Tetramin (Tetra, Melle, Germany), supplemented with 48 hours live brine shrimp (*Artemia* spp) one week before the onset of reproduction. During the bioassay the amount of food delivered was adjusted according to fish development and size. The experiment consisted of six treatments in duplicate: a control (dechlorinated water); a solvent control (0.0002% acetone, ACET), and four SIM treatments, 8, 40, 200 and 1000 ng/L concentrations. The selection of SIM treatments was based in previous studies (Neuparth et al., 2014), taking into consideration environmentally relevant concentrations. SIM (CAS no. 79902-63-9; Sigma Aldrich) was dilute in ACET to obtain a stock solution of 4 mg/mL. From this solution, working solutions were prepared by serial dilutions of the stock solution and aliquots were stored in the dark at −20°C until use. All solutions were prepared in order to have a final ACET concentration of 0.0002%. Based on preliminary tests, in order to maintain exposure concentrations, the working solutions were dosed directly into the water twice a day; in the morning (T _Oh_) and in the afternoon (T _8h_), in a volume that was equivalent to the water renewal during that period.

### 2.4. Reproductive capability

Reproductive capability studies were performed 70 days after the beginning of the chronic exposure for all the tested groups (120 days age-old fish). Reproductive success was assessed through the evaluation of two endpoints: fecundity (number of eggs per female per day) and percentage of fertilization (% of fertilized eggs per female per day). In the afternoon, before the beginning of the trials, each aquarium was divided in two separated section in which a suspended cage with a net bottom covered with marbles was fixed. Males and females were equally distributed through the cages in a manner that matched the sex ratio of the respective treatment (four sub-replicates per treatment) (Figure 1.A). The sex ratio was determined by visual inspection of each animal and confirmed at the end of the bioassay by stereomicroscope gonads observation.

**Figure 1.**
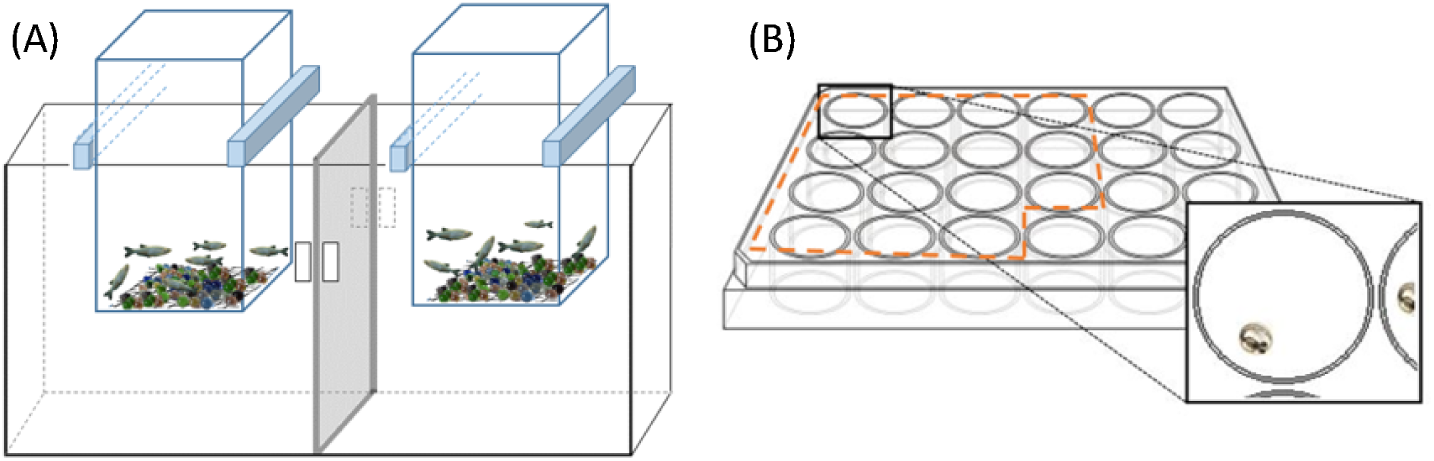
Schematic representations of the breeding setup for reproductive capability assay (A) and distribution of the embryos in the 24 well plate for embryogenesis studies (B).

During five consecutive days, 1-1.5 hour after the beginning of the light period, breeding fish were removed and the eggs collected, cleaned and conserved in 70% ethanol, for posterior counting and determination of the percentage of fertilized eggs.

### 2.5. Embryogenesis studies

Embryogenesis studies were carried out with slight modification of OECD TG236 (OECD, 2013 and Torres et al. (2016). During the reproductive trials, 15 embryos from each treatment sub-replicate, with 1-1.5 hpf, were kept and randomly placed in 24 well plates (one egg per well) with 2 ml of clean dechlorinated water, free of SIM (Figure 1.B). The 24 well plates were randomly maintained on a water bath at 26.5 ± 0.5°C during 80 h. Embryos were checked, every day, for mortality and at the end of the assay (80 hpf) were observed, under a stereomicroscope, for embryo development analysis. Morphological abnormalities on eyes, head, tail or yolk-sac; pericardial oedema; abnormal cell growth and developmental arrest were recorded as present or absent (Torres et al., 2016). Heart rate was evaluated during 15 s in four embryos per plate using a stop-watch, restarting the counting if the embryo moved.

### 2.6. Sampling

At end of the assay, the animals were sacrificed with an anaesthetic overdose of 300 mg/L tricane methanesulfonate (MS-222), to which sodium bicarbonate was added to prevent the acidification of the solution. All animals were measured and weighted, and Fulton’s condition factor (K) was determined (K=(weight / lenght^3^)×100) – Nash et al., 2006). Livers were collected and i) preserved in RNA Later for gene expression analysis, ii) frozen in liquid nitrogen and stored at −80 °C for further cholesterol and triglycerides quantification.

### 2.7. Lipid extraction and cholesterol and triglycerides quantification

Lipids were extracted from the liver samples using a low toxicity solvent extraction protocol, adapted from Schwartz and Wolins (2007). The tissues were homogenized in 10 mM PBS buffer pH 7.4 containing 10 mM EDTA (10 mg of tissue per 1 mL buffer with two ceramic beads) on a Precellys 24 homogenizer. 500 µl of homogenates were then transferred, in duplicate, to test glass vials containing 5 mL of isopropanol/hexane solution (4:1). The samples were vortexed for 1 min and incubated at room temperature in the dark with constant shaking for 2 h. In order to avoid lipid peroxidation, vials were always passed through nitrogen current before closing. After the incubation period, samples were washed with 2 mL of petroleum ether/hexane solution (1:1). Vials were again vortexed for 1 min and left in the dark at room temperature for 10 min. The phases were then separated by adding 1 mL of Milli-Q water, vortexed for 1 min, incubated in the dark at room temperature, with constant shaking for 20 min, and centrifuged at 1000 × G, for 10 min. The upper phase containing the lipids was collected into new vials and evaporated to dryness under nitrogen current. Dried extracts were then stored at −20 °C until cholesterol and triglycerides quantification.

Dry extracts were re-suspended in 100 µL of isopropanol and sonicated in ultra-sound bath Bandelin Sonorex RK100H for 15 min at room temperature. Quantification of cholesterol and triglycerides was performed through enzymatically colorimetric assays using Infinity Cholesterol Liquid Stable Reagent and Infinity Triglycerides Liquid Stable Reagent, respectively, both purchased from Thermo scientiphic, Biognóstica and following the manufacturer’s protocols. Samples were measured in duplicate and absorbance determined at λ 490 nm using a microplate reader (Biotech Synergy HT) coupled with software Gen5 (version 2.0). In every run, a standard curve was performed for the cholesterol and triglycerides optical quantification. Cholesterol and Triolin standards were prepared and subjected to 6 serial dilutions (from 0.156251 to 5 mg/mL for cholesterol and 0.0625 to 2 mg/mL for triolin).

### 2.8. Gene expression

#### 2.8.1. RNA isolation and cDNA synthesis

Liver samples from the different treatments (approximately 3 mg) were used to isolate total RNA via the Illustra RNAspin Min RNA Isolation Kit (GE Healthcare), according to the manufacturers’ protocol. RNA quantification was performed on a Take3 Micro-Volume Plate Reader (Biotech Synergy HT) coupled with the software Gen5 (version 2.0). RNA quality was verified by electrophoresis in 1.5 % agarose gel and through the measurement of the ratio of absorbance at λ 260 / λ 280 nm. Total cDNA was generated from 1 µg of total RNA extracted using the iScriptTM cDNA Synthesis Kit (Bio-Rad).

#### 2.8.2. qRT-PCR

Fluorescence-base quantitative real time PCR (qRT-PCR) was used to evaluate the transcription profiles of several genes involved in the mevalonate pathway with important roles in the cholesterol biosynthesis. The 3-hydroxy-3-methylglutaryl-CoA reductase (*hmgcr*) was selected because it is the primary target of SIM being the limiting step in the mevalonate pathway. According to previous studies on the HMGCR tissue distribution, we chose to assess *hmgcra* instead of *hmgcrb*, due to its predominance in the liver, while *hmgcrb* prevails in the brain (Al-Habsi et al., 2016). Other two genes, lanosterol 14α-demethylase (*cyp51*) and 7-dehydrocholesterol reductase (*dhcr7*) were chosen as intermediate genes in the mevalonate pathway. Ribosomal protein L8 (*rpl8*) was used as reference gene, as its expression levels remained constant across treatments. All the primers were already described by other authors (Table 1). For each treatment (n = 8), the expression of individual target genes was performed with the Mastercycle ep realplex system (Eppendorf). cDNA of liver samples was amplified in duplicated in 96-well optical plates, containing 10 µL of NZY qPCR Green Master Mix (2x) (NZYTech), 0.8 µL of each primer (forward and reverse), 2 µL of cDNA at 100 nmol and 6.4 µL of water in order to reach a final reaction volume of 20 µL. On each plate, a non-template control was included. In order to determine the efficiency of the reaction, a two-step qRT-PCR was performed: initial denaturation at 95 °C (3 min), followed by 40 cycles of amplification with a denaturation at 95 °C for (15 s) and combined annealing and extension at 58 – 62°C, depending on the pair of primers (25 s) (Table 1). A melting curve (from 55 °C to 95 °C) was generated in each run to confirm the specificity of the reactions. The PCR products were analysed by gel electrophoresis in 2 % agarose to check the presence of single bands with the expected size between 136 and 199 bp, depending on the pair of primers (Table 1). The PCR efficiency for the genes of interest and for the reference gene was determined by a standard curve, using six serial dilutions of cDNA pools of all samples (from 0.064 to 200 ng of cDNA). The minimum efficiency obtained was 94% (Table 1). Relative change in the transcription abundance of target genes was normalized to *rpl8* and calculated using the Livak method (Livak and Schmittgen, 2001). Control expression levels (ACET treatment) were normalized to 1 and data were then expressed as fold changes relative to solvent control group.

**Table 1.**
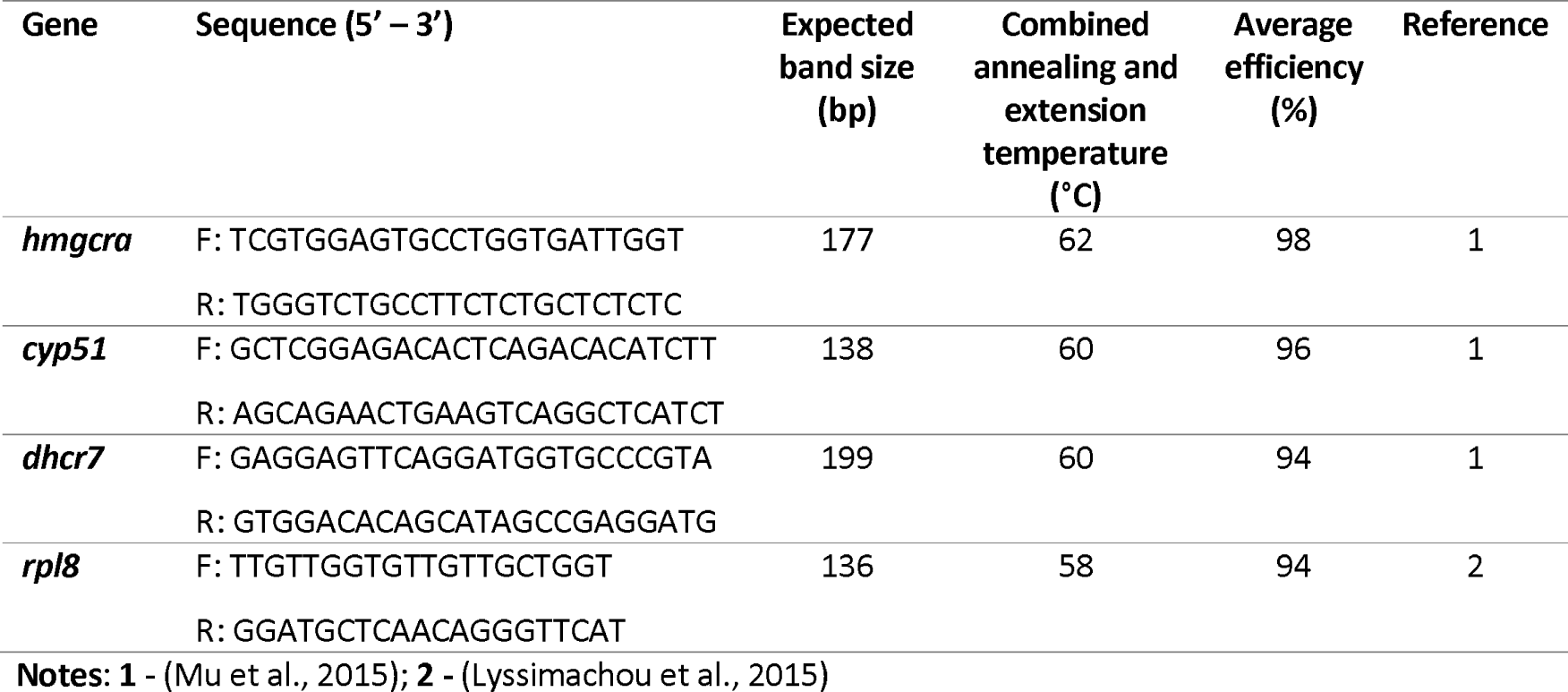
Primers, forward (F) and reversed (R), and parameters used in the qRT-PCR for gene expression quantification in the liver of *D. rerio.*

### 2.9. Simvastatin analytic quantification by liquid chromatography – tandem mass spectrometry

The actual concentrations of SIM were determined, in each treatment, twice during the bioassay: 30 min after the first SIM spike of the day (T_0h_) and 8 hours after the first SIM spike and immediately before the second spike (T_8h_). Two samples from each treatment (bulk samples from each replicates) were collected and stored at −20 °C for posterior quantification by solid-phase extraction (SPE) and Liquid Chromatography – Tandem Mass Spectrometry (LC-MS/MS). One hundred millilitres of each water sample were filtered through 0.45µm PVDF filters (Millipore) and spiked with 50 ng/L of Mevastatin (MEV, Sigma-Aldrich) as internal standard. Subsequently, the sample was loaded into a 60 mg Oasis HLB (Waters) cartridge previously conditioned with 5 mL of methanol (MeOH) and 5 mL of Milli-Q water. The cartridges were washed with 5 mL of Milli-Q water, dried under a N_2_ stream and eluted with 5 mL of MeOH. The final extracts were evaporated to dryness under N_2_ stream, reconstituted using 100 µL of MeOH and injected in the LC-MS/MS system. The quantification was performed by the matrix-matched method, by spiking with SIM real water in the 5 to 1000 ng/L range and submitting these samples to the entire protocol. Due to the fast conversion of SIM and MEV to their corresponding hydroxyacid forms (SIMHA and MEVHA) in aqueous media (Bews at al., 2014), these latter forms were also quantified and the final concentration was calculated as sum of the lactone and hydroxyacid forms.

The LC–MS/MS system (Varian) was equipped with two ProStar 212 high-pressure mixing pumps, a vacuum membrane degasser, an autosampler and a thermostated column compartment ProStar 410 module (Varian). A volume of 10 µL of extract was directly injected into a Phenomenex Luna C18 (50 mm × 2.1 mm, 3 μm particle diameter) maintained at a constant temperature of 30 °C. The target compounds were separated at a flow rate of 0.2 mL min^−1^ using 0.1% of formic acid in both, Milli-Q water (A) and MeOH (B) as eluents. The applied gradient was as follows: 0–1 min, 30% B; 1–15 min, linear gradient to 100% B; 15–18 min 100% B; 18–18.5 min, linear gradient to 30%B and 18.5–23 min, 30% B. The system was interfaced to a Varian 320-MS triple quadrupole mass spectrometer equipped with an electrospray interface. Nitrogen was used as nebulizing and drying gas and Argon was used as collision gas. The analytes were determined in the electrospray positive (SIM and MEV) and negative (SIMHA and MEVHA) and multiple-reaction monitoring (MRM) mode of acquisition. Two MRM transitions were used for each compound quantifier and qualifier respectively (precursor > product ion, m/z values): 441 > 325 and 419 > 199 for SIM, 413 > 311 and 391 > 185 for MEV, 435 > 319 and 435 > 341 for SIMHA and finally, 407 > 305 and 407 > 327 for MEVHA.

The method limits of quantification (LOQs) were 1.5 and 1.2 ng/L for SIM and SIMHA, respectively. The linearity, evaluated in terms of correlation coefficient for the sum of SIM and SIMHA, was R^2^ 0.9964 and the recovery was 91%.

### 2.10. Statistical analysis

The obtained data were checked for homogeneity of variances (Leven’s test) and normality (Kolmogorov-Smirnov test) and subsequently analysed by one-way ANOVA. Post-hoc comparisons were carried out using Fisher’s least significant difference (LSD) test. When differences were not found between control and solvent control groups, these two treatments were pooled and referred to as control. Significant differences were set as p < 0.05. All the statistics were computed with Statistica 13 (Statsoft, USA).

## 3. Results

### 3.1. Simvastatin analytical quantification

Table 2 summarizes the actual SIM concentrations measured during the bioassay at times T_0h_ and T_8h_. Due to the high conversion of SIM to SIMHA in aqueous media (Bews et al. 2014), the final concentration was calculated as sum of both active forms and given as SIM. No SIM was detected in solvent control group. Results show that SIM concentrations at the initial time (T_0h_), were marginally lower than nominal ones. At T_8h_, the SIM concentration decays as expected, but was still present in all treatments (except control) with a minimum and maximum decay of 27.9% and 57.3% in the 8 and 1000 ng/L SIM treatments, respectively.. To ensure the exposure of zebrafish to concentration close to the nominal throughout the assay, the aquaria were spiked with SIM twice a day.

**Table 2.** Nominal and measured concentrations of SIM in water samples collected in duplicate from each treatment after the first contamination of the day (T _Oh_)and immediately before the second one (T _8h_), Data are expressed as mean ± standard error.

| Time | Solvent control | SIM 8 ng/L<sup>a</sup> | SIM 40 ng/L<sup>a</sup> | SIM 200 ng/L<sup>a</sup> | SIM 1000 ng/L<sup>a</sup> |
| --- | --- | --- | --- | --- | --- |
| T 0 h | n.d.<sup>b</sup> | 6.8 ± 0.45 | 37.9 ± 3.21 | 108.6 ± 3.70 | 751.7 ± 40.08 |
| T 8 h | n.d.<sup>b</sup> | 4.9 ± 0.08 | 23.0 ± 0.08 | 74.0 ± 1.47 | 343.4 ± 23.14 |

<sup>a</sup> Nominal SIM concentrations
<sup>b</sup> Not detected

### 3.2. Survival, growth, body weight and length

Table 3 displays the survival rates, weight and length of zebrafish males and females following the 90-days chronic exposure to environmentally relevant SIM concentrations. No significant differences among treatments were observed in the survival rates; almost 100 % of the fish were alive at the end of the bioassay. In SIM exposed fish, a significant decrease of 6.14 % in the body weight was observed in males exposed to the highest SIM concentration (1000 ng/L), compared to the control groups. In contrast, the weight of females, the length and the condition factor of both sexes were not affected by SIM exposure.

**Table 3.**
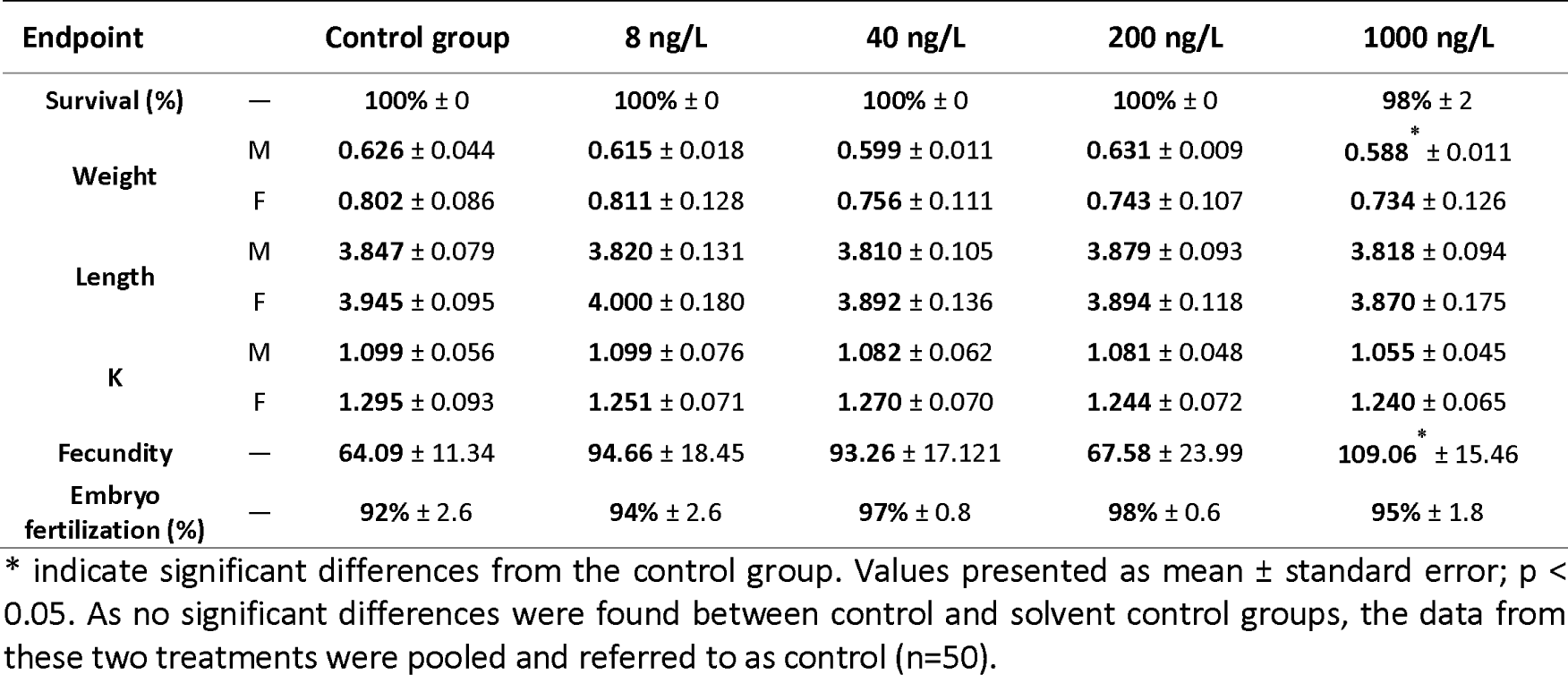
Chronic effects of SIM on survival, weight, length, Fulton’s condition factor (K), fecundity (number of embryos/female/day), and % of fertilization (% of fertilized eggs/female/day) of *D. rerio* after 90 days of exposure (M, males; F, females).

### 3.3. Reproductive capability

Females from all tested SIM concentrations exhibited a trend towards an increase in fecundity when compared with females from control group (Table 3), reaching statistically significance at the 1000 ng/L SIM treatment. This increase was not followed by the percentage of fertilization, for which significant differences were not found.

### 3.4. Embryogenesis

The cumulative rate of embryos mortality at 80 hpf was similar among treatments and kept at low levels, that ranged from 5 % (200 ng/L SIM parental exposure) to 11.7 % (1000 ng/L SIM parental exposure) (figure 2. A). From all the analysed abnormalities, tails anomalies (shortened and/or curled tails – Figure 2. B) were the ones recorded in higher number in SIM exposed groups. A significant increase of tail anomalies was observed at the lowest SIM concentrations, i.e. 10-fold increase in both 8 and 40 ng/L, compared with the control group. A significant increase of total abnormalities (sum of head, tail, eyes and yolk-sac anomalies plus pericardial oedema) was also observed in embryos from the lowest SIM concentration (8 ng/L), with 18.3 % of abnormal embryos in comparison with 0.83 % of the control group (figure 2. C). Similarly, 15 % of abnormal embryos was observed at 40 ng/L SIM, although significance was not reached (p=0.13). Furthermore, SIM parental exposure was also able to significantly increase heart beat by 8.37 bpm at 40 ng/L exposure treatment relative to control group (Figure 2. D).

**Figure 2.**
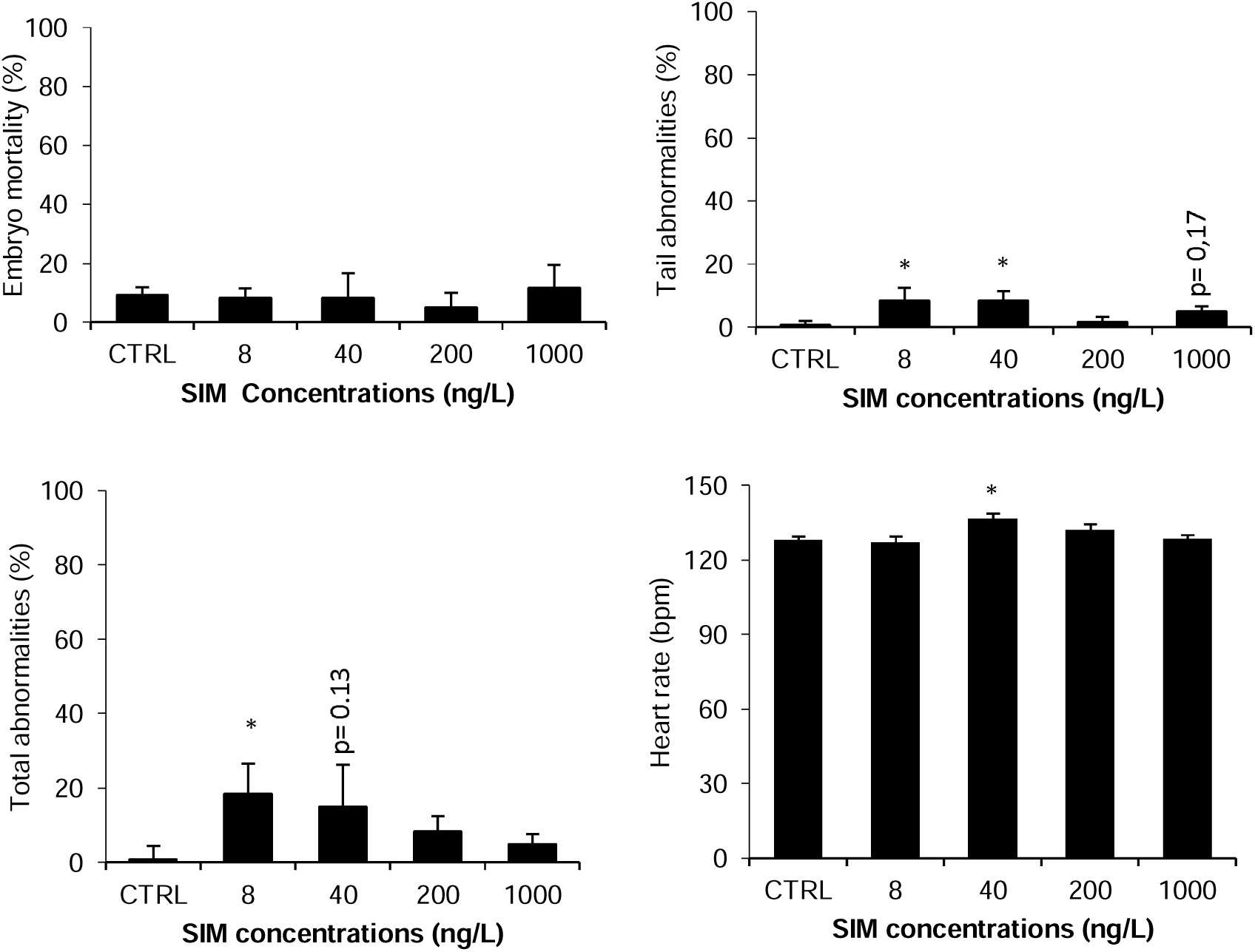
Cumulative mortality – n=60 (A); Anomalies in the tail – n=60 (B); Total abnormalities – n=60 (C); and Heart rate – n=32 (D) observed in zebrafish embryos at 80 hpf, following parental chronic exposure to SIM. Error bars indicate standard errors; * indicate significant differences from the control group (p < 0.05). As no significant differences were found between control and solvent control groups, the data from these two treatments were pooled and referred to as control (CTRL).

### 3.5. Hepatic cholesterol and triglycerides content

SIM significantly decreased liver cholesterol (Chol) levels in female zebrafish at intermediate SIM concentrations, i.e. 40 and 200 ng/L, with a 53.56 and 64.59 % (p=0.01) decrease, respectively, in comparison with the control group (Figure 3. A). Males exhibited the same Chol decrease pattern, although significance was not reached. Both males and females exhibited a slight Chol increase in the highest SIM concentration (1000 ng/L) when compared with the intermediate SIM treatments, nevertheless the Chol levels remained lower than the ones from the control group. Chol levels presented a non-monotonic dose response curve with a decrease at intermediate SIM concentrations (40 and 200 ng/L).

**Figure 3.**
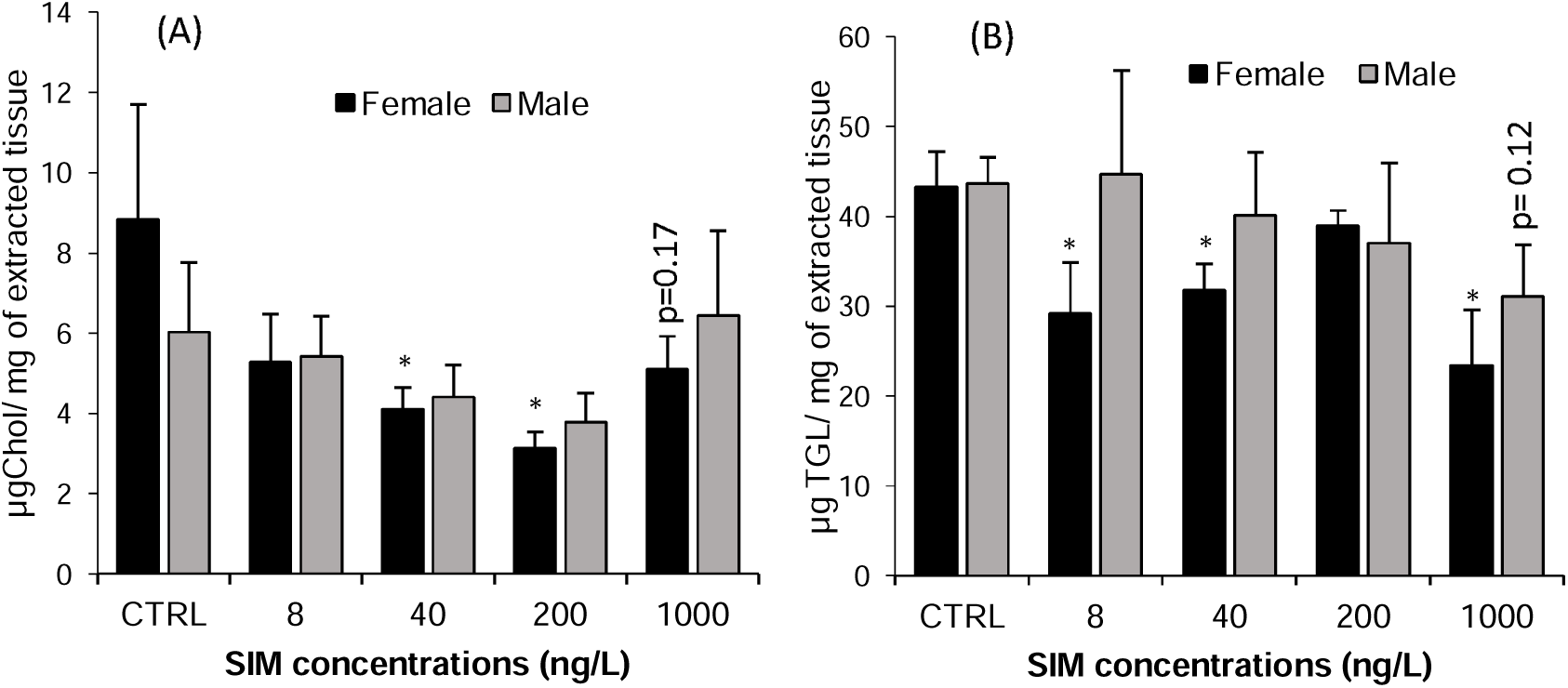
Chronic effects of SIM on *D. rerio* liver cholesterol (A) and triglycerides (B) content, after 90 days of exposure. Error bars indicate standard errors; * indicate significant differences from the control group (p < 0.05). As no significant differences were found between control and solvent control groups, the data from these two treatments were pooled and referred to as control (n=8).

Triglycerides (TGL) levels in female liver were significantly lower in all SIM concentrations, when compared with control group, with the exception of the 200 ng/L treatment (Figure 3. B). Exposure to 8, 40, and 1000 ng/L of SIM was able to significantly decrease TGL levels by 32.59 (p=0.02), 26.57 and 45.89 % (p= 0.0017), respectively. No significant differences in TGL levels were observed in males.

### 3.6. Gene expression

Both *D. rerio* females and males showed significant differences in the transcription levels of several genes of the mevalonate pathway, after 90 days of exposure to SIM (Figure 4). Similar to Chol levels, *hmgcra* e *cyp51* genes presented, for both sexes, a non-monotonic dose-response curve with a likewise decrease at 40 and 200 ng/L SIM. In females, *hmgcra* was significantly downregulated by 2.64 (p= 0.01) and 2.89-fold (p= 0.01) in comparison with the solvent control treatment (ACET) after exposure to 40 and 200 ng/L of SIM, respectively. Similarly, in males, a 4.86 (p= 0.02) and 2.94-fold significant decrease in *hmgcra* expression levels was observed at the same treatments, (40 and 200 ng/L SIM, respectively). In females, none of the remaining genes showed altered transcript levels. Nevertheless, a trend towards a decrease in *cyp51* gene expression was observed at 40 ng/L SIM, although significance was not reached (p = 0.07). Males, on the other hand, exhibited a 6.80-fold significant decrease of *cyp51* expression when exposed to 40 ng/L SIM. A similar decrease pattern was observed at 200 ng/L SIM, just in the range of significance (p=0.06). Neither males nor females presented significant alterations in *dhcr7* transcription levels. No significant differences were observed between control and solvent control for the analysed genes, with the exception of the *dhcr7* in males.

**Figure 4.**
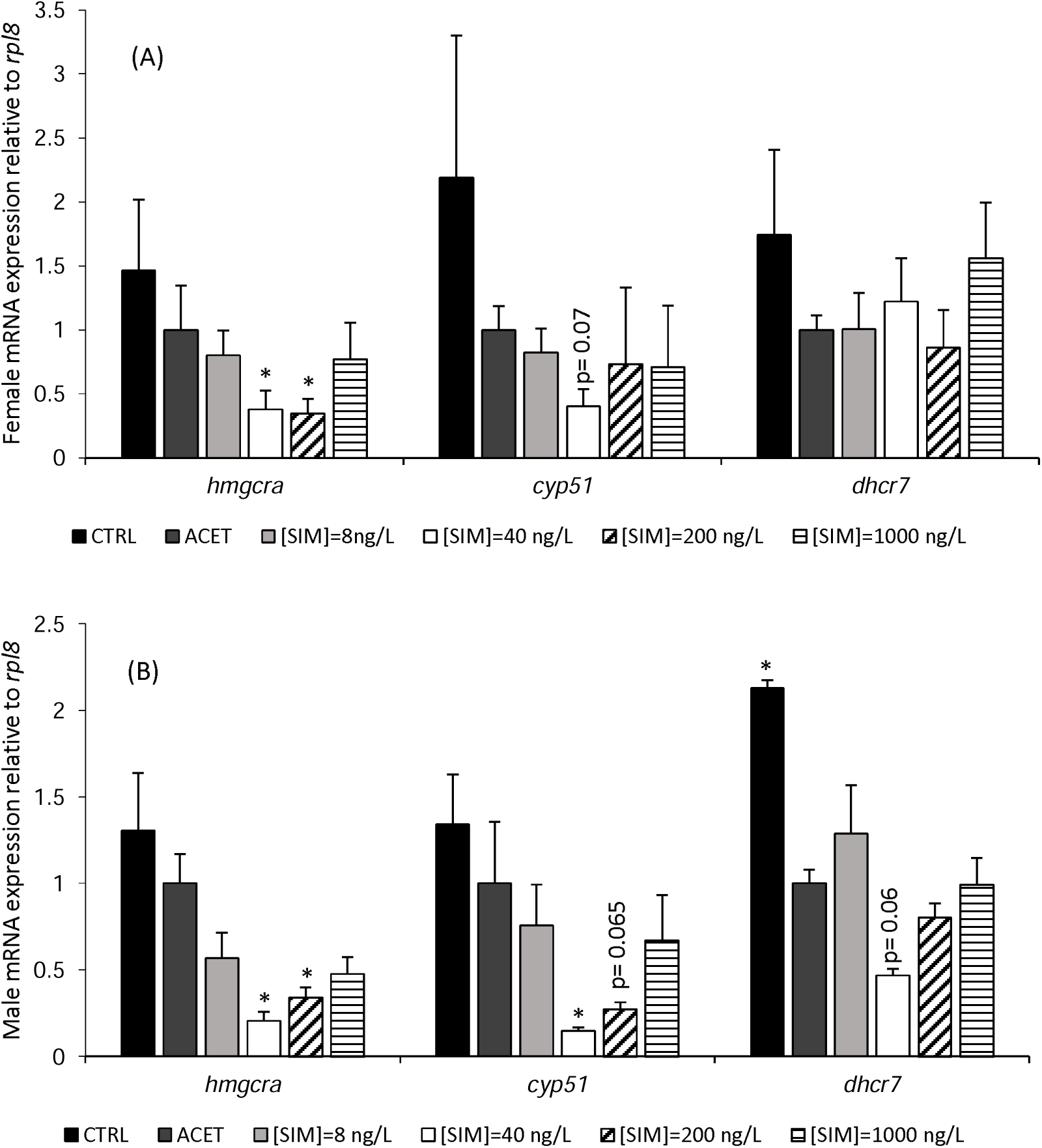
Females (A) and Males (B) relative mRNA expression of *hmgcra, cyp51*, and *dhcr7* in adult *D. rerio* livers after 90 days SIM exposure. Error bars indicate standard errors; asterisks (*) indicate significant differences from the solvent control treatment (ACET = 1-fold) (p< 0.05) (n=8).

## 3. Discussion

Statins, such as SIM, are designed to produce in humans specific biological effects at low concentrations and, therefore, once in the aquatic environment many aquatic taxa might be at risk. Despite the ubiquitous nature of statins in the aquatic ecosystems and the liability of the scientific community to improve their risk assessment, the chronic effects of these pharmaceuticals in aquatic organisms has not yet been adequately studied. In order to address this actual gap of knowledge, the present study aimed at investigating the chronic partial life-cycle and early offspring embryonic development effects of SIM exposure, using zebrafish as model organism. Thus, several fundamental questions were addressed: Does chronic SIM exposure affect ecological relevant endpoints in zebrafish? Are these effects related with alteration of cholesterol and triglycerides levels? Is the cholesterol homeostasis disturbed and correlated with changes in the transcription of key genes from the mevalonate pathway?

In mammals, SIM and other statins operate by competitively inhibiting the rate limiting enzyme of the mevalonate pathway (HMGCR), thereby reducing Chol biosynthesis (Blumenthal, 2000; Endo et al., 1976a; Endo et al., 1976b; Sehayek et al., 1994). Given the conservation status of the mevalonate pathway among mammals and fish and the high degree of conservation of HMGCR, with respect to its catalytic activity and statin interactions across all metazoans (Santos et al., 2016), it is expected that fish display similar responses to those described in mammals, i.e. an impact on Chol synthesis and alterations in the transcription of genes from the mevalonate pathway. The results of this study show that chronic exposure to low levels of SIM impacts key ecologically-relevant endpoints of zebrafish, particularly embryonic development, concomitantly with alterations in the Chol and TGL levels in the liver. Furthermore, an impact in the transcription of key genes from the mevalonate pathway was also observed. Importantly, several of the analysed endpoints did not follow a dose-response curve, with significant effects observed only in the low-intermediate SIM concentrations. The results also indicate for some endpoint sex-dependent differences, suggesting that females and males could respond differently to SIM exposure.

The sub-lethal exposure to the highest SIM dose (1000 ng/L) significantly decreased males’ body weight. To our knowledge, there is no scientific study reporting SIM effects on the weight control of fish or mammals and this alteration was not reflected in any of the biological endpoints analysed, in the levels of Chol/TGL, nor in the transcription level of the screened genes. Therefore, this SIM effect might be related with the intrinsic toxicity of this pharmaceutical. On the other hand, females’ exposure to SIM resulted in an increase in fecundity that reached significance at the higher SIM concentration (1000 ng/L). We hypothesize that a hormesis phenomenon could have been responsible for the unexpected stimulating effect of SIM in zebrafish fecundity. Hormesis has been documented in a wide range of vertebrates and invertebrates in response to a variety of stressors, being characterized as a stimulatory effect on biological parameters, particularly reproduction and/or growth, in response to low-doses of toxicants (Ayyanath et al., 2015; Calabrese, 1999; Correia, et al., 2001; Guo & Chen, 2015; Lefcort et al., 2008; Meador et al., 2011; Stebbing, 1982). This is view as an adaptive response that overcompensates the homeostatic regulatory control mechanisms induced by low levels of stress or damage, which will therefore be responsible to enhance the activities of the physiological systems (Calabrese & Baldwin, 2001).

Our results on the embryonic development showed that chronic parental exposure to 8 and 40 ng/L SIM resulted in a significant increase of morphologic abnormalities, being the tail anomalies the most frequent abnormalities observed. Also, an increase in the heart rate at 40 ng/L SIM parental exposure was observed. Some previous studies that tested higher SIM concentrations showed that the embryonic development of zebrafish is a target of SIM with described embryo malformations similar to those reported here in addiction to abnormalities in eyes and yolk-sac, pericardial oedema and impairment of primordial germ cell migration (Campos et al., 2016; Cunha et al., 2016; Ribeiro et al., 2015; Thorpe et al., 2004). It is plausible to hypothesize that the parental exposure might be responsible for the significant embryonic abnormalities observed. Thus, the observed disruption in the zebrafish embryonic development induced by SIM could have important ecological impacts at the fish population level, affecting their fitness and survival, since the impact of SIM not only persist in the exposed parents but also can be passed to the second generation as the embryos were raised in SIM-free water.

The findings recorded on the embryonic development integrate well with the observed decrease of Chol and TGL levels in females’ liver exposed to the intermediate SIM concentrations. In fact, the most abundant lipids in vertebrate embryos (including zebrafish) are Chol and TGL, which are transferred from the mother to the embryo yolk, having an important role in providing energy and cellular structural components during the embryogenesis (Fraher et al., 2016). On the other hand, the study of Cunha et al. (2016) reported that zebrafish embryos exposed to 5 and 50 µg/l of SIM presented a transcriptional downregulation of the retinoic acid receptor (RAR) and retinoid X receptor (RXR), which have been implicated in lipid homeostasis (RXR) and development (RXR and RAR) (Castro and Santos, 2014). Overall, our findings and the ones from Cunha et al. (2016) point to an impact of SIM in lipid homeostasis in zebrafish embryos that might be implicated in the observed abnormalities in the embryogenesis.

Chol and TGL are also essential in fish adulthood for the maintenance of several biological processes like growth, reproduction and cell integrity (van der Wulp et al., 2013; Vergauwen et al., 2010). The administration of statin, leads to a reduction of those lipids, at least in mammals. Although the most prominent effects attributable to statins are their potent Chol lowering properties by the inhibition of HMGCR enzyme, it is also well established that statins significantly reduce TGL levels (Funatsu et al., 2001). The reduction of TGL levels by statins seems to be related with the scarce secretion of very low-density lipoprotein (VLDL) from the liver and the increase in clearance of triglyceride-rich lipoprotein via inducted LDL receptors from plasma (Funatsu et al., 2001). As expected, by the studies with mammals, our results show that chronic low levels exposure to SIM significantly reduce the Chol and TGL content in zebrafish. Interestingly, SIM treatment was more effective in females than males with significant differences registered for both lipids only in females at the intermediate SIM concentrations. Although not exhibiting significant decreases in Chol and TGL levels at the SIM treatments tested, males present similar response pattern to that of females, at least for Chol levels. Similar sexual dimorphism on Chol and TGL levels have been reported by Al-Habsi et al. (2016) after feeding zebrafish with atorvastatin for 30 days; registering a significant reduction of both Chol and TGL only for females. Altogether, these suggest that female zebrafish are more prone to the effects of SIM on Chol and TGL content than males. In fact, female zebrafish need higher lipid content for reproduction than males due to the importance of these components in egg production (Landgraf et al., 2017).

This study shows that the observed SIM-induced biological and biochemical effects occurred in parallel, with significant alterations in gene transcription in male and females’ liver of two key mevalonate pathway genes, *hmgcra* (the rate limiting step in the mevalonate pathway) and *cyp51* (an intermediate gene in the mevalonate pathway). A downregulation of these two genes was observed in males and females at the intermediate concentrations of SIM (40 and 200 ng/L for *hmgcra* and 40ng/L for *cyp51*). Interestingly, these are the SIM concentrations that significantly reduced the levels of liver Chol and therefore, the observed changes in transcript levels of key genes of the mevalonate pathway could be linked with the alterations in Chol levels. The mechanism of SIM here observed is similar to the reported in mammals, i.e. decrease of Chol production concomitantly with changes in the transcription levels of gene encoding the enzyme responsible for the limiting step of mevalonate pathway – HMGCR. However, the majority of the studies available in the literature, for humans and mammalian animal models, report a transcriptional up-regulation of *hmgcr* after SIM administration (Conde et al., 1999; Gouni-Berthold et al., 2008; Rudling et al., 2002). Also, one study with zebrafish fed with the statin, atrovastin, for 30 days presented the same *hmgcr* pattern to that reported in mammals (Al-Habsi et al., 2016), in contrast with the downregulation observed in the present study. The contrasting expression profile of *hmgcr* observed in the present study suggests that SIM affects zebrafish transcription levels of *hmgcr* in a time-specific manner. To the best of our knowledge, there aren’t studies available that investigate the time effects of SIM on gene expression of *hmgcr*, however, a clear time-dependent gene expression response is descried in the literature for distinct genes in several aquatic model organisms exposed to different compounds, including SIM (Cunha et al., 2016; Hamadeh et al., 2002; Kim et al., 2015). Thus, the absence of data on the temporal *hmgcr* gene expression highlights the importance of using multiple time-points in gene expression studies with statins. It is worth noting that SIM induced significant toxic responses at the lowest concentrations (8, 40 and/or 200 ng/L) in most of the analyzed endpoints, while at the highest concentration (1000 ng/L), the responses were similar to the control conditions. Therefore, the intermediate SIM exposures appeared more effective to disrupt homeostasis than the higher exposure. These responses appeared to be non-monotonic, showing an inverted U-shape for the embryonic development abnormalities and a U-shape for the cholesterol levels and for the expression of *hmgcra* and *cyp51* genes. This is a common pattern observed in low concentration exposures during chronic experiments for some endocrine disrupting chemicals; therefore the non-monotonicity responses should not be ignored in environmental studies, neither in hazard assessment (Vandenberg et al., 2012). The mechanism(s) that cause the non-monotonicity of SIM in the present study are currently unknown and should be investigated. However, we hypothesize that the interactions of the endogenous feedback mechanism to maintain the homeostatic control of Chol and the mechanisms of SIM to reduce the Chol levels might have a role in the non-monotonic responses here reported. The lowest SIM exposure may result in the activation of SIM associated-pathways to reduce the Chol levels, however, at the higher doses, the endogenous mechanisms to control the levels of Chol must have compensated the effects of SIM in order to achieve the homeostasis and the effect of SIM was less evident.

In summary, taken together the findings of the present study, low environmentally realistic concentrations of SIM have a significant impact in zebrafish Chol metabolism with some similarities to those reported in mammals. Chronic exposure to SIM reduced the levels of Chol and TGL, mainly in females, concomitantly with changes in the transcription levels of key genes involved in the Chol biosynthesis. These alterations could be associated with the disruption of embryonic development, which might have implications at an ecological level, given the effects observed in the low ng/L range. Given that SIM is expected to impact the mevalonate pathway in a disparate range of animal taxa (Santos et al., 2016), the results of the present study highlight the need to expand chronic low-level exposure studies to other groups of animals.

## 4. Acknowledgements

T. Neuparth was supported by the Postdoctoral fellowship SFRH/BPD/77912/2011 from Foundation of Science and Technology, Portugal. R. Montes, R. Rodil and J.B. Quintana acknowledge financial support from the Spanish “Agencia Estatal de Investigación” (project no. CTM2014-56628-C3-2-R), the Galician Council of Culture, Education and Universities (ED431C2017/36) and FEDER/ERDF.

## References

Al-Habsi, A. A., Massarsky, A., & Moon, T. W. (2016). Exposure to gemfibrozil and atorvastatin affects cholesterol metabolism and steroid production in zebrafish (Danio rerio). Comparative Biochemistry and Physiology Part – B: Biochemistry and Molecular Biology, 199, 87–96. https://doi.org/10.1016/j.cbpb.2015.11.009

Arnold, K. E., Brown, A. R., Ankley, G. T., & Sumpter, J. P. (2014). Medicating the environment: assessing risks of pharmaceuticals to wildlife and ecosystems. Philosophical Transactions of the Royal Society of London. Series B, Biological Sciences, 369, 20130569. https://doi.org/10.1098/rstb.2013.0569

Ayyanath, M. M., Scott-Dupree, C. D., & Cutler, G. C. (2015). Effect of low doses of precocene on reproduction and gene expression in green peach aphid. Chemosphere, 128, 245–251. https://doi.org/10.1016/j.chemosphere.2015.01.061

Azzouz, A., & Ballesteros, E. (2012). Combined microwave-assisted extraction and continuous solid-phase extraction prior to gas chromatography-mass spectrometry determination of pharmaceuticals, personal care products and hormones in soils, sediments and sludge. Science of the Total Environment, 419, 208–215. https://doi.org/10.1016/j.scitotenv.2011.12.058

BIO Intelligence Service (2013), Study on the environmental risks of medicinal products, Final Report prepared for Executive Agency for Health and Consumers

Blumenthal, R. S. (2000). Statins: Effective antiatherosclerotic therapy. American Heart Journal, 139, 577–583. https://doi.org/10.1016/S0002-8703(00)90033-4

Burg, J. S., & Espenshade, P. J. (2011). Regulation of HMG-CoA reductase in mammals and yeast. Progress in Lipid Research, 50, 403–410. https://doi.org/10.1016/j.plipres.2011.07.002

Calabrese, E. J. (1999). Evidence that hormesis represents an “overcompensation” response to a disruption in homeostasis. Ecotoxicology and Environmental Safety, 42, 135–137. https://doi.org/10.1006/eesa.1998.1729

Calabrese, E. J., & Baldwin, L. A. (2001). Hormesis: U-shaped dose responses and their centrality in toxicology. Trends in Pharmacological Sciences, 22, 285–291. https://doi.org/10.1016/S0165-6147(00)01719-3

Campos, L. M., Rios, E. A., Guapyassu, L., Midlej, V., Atella, G. C., Herculano-houzel, S., Benchimol, M., Mermelstein, C., Costa, M. L. (2016). Alterations in zebrafish development induced by simvastatin1: Comprehensive morphological and physiological study, focusing on muscle. Experimental Biology and Medicine, 241, 1950–1960. https://doi.org/10.1177/1535370216659944

Conde, K., Roy, S., Freake, H. C., Newton, R. S., & Fernandez, M. L. (1999). Atorvastatin and simvastatin have distinct effects on hydroxy methylglutaryl-CoA reductase activity and mRNA abundance in the guinea pig. Lipids, 34, 1327–1332. https://doi.org/10.1007/s11745-999-0485-2

Correia, A. D., Costa, F. O., Neuparth, T., Diniz, M. E., & Costa, M. H. (2001). Sub-lethal Effects of Copper-Spiked Sediments on the Marine Amphipod Gammarus locusta1: Evidence of Hormesis1? Ecotoxicology and Environmental Restoration, 4, 32–38.

Craig, P. M., & Moon, T. W. (2011). Fasted Zebrafish Mimic Genetic and Physiological Responses in Mammals: A Model for Obesity and Diabetes? Zebrafish, 8, 109–117. https://doi.org/10.1089/zeb.2011.0702

Cunha, V., Santos, M. M., & Ferreira, M. (2016). Simvastatin effects on detoxification mechanisms in Danio rerio embryos. Environmental Science and Pollution Research, 23, 10615–10629. https://doi.org/10.1007/s11356-016-6547-y

Dahl, U., Gorokhova, E., & Breitholtz, M. (2006). Application of growth-related sublethal endpoints in ecotoxicological assessments using a harpacticoid copepod. Aquatic Toxicology, 77, 433–438. https://doi.org/10.1016/j.aquatox.2006.01.014

Daughton, C. G. (2016). Pharmaceuticals and the Environment (PiE): Evolution and impact of the published literature revealed by bibliometric analysis. Science of the Total Environment, 562, 391–426. https://doi.org/10.1016/j.scitotenv.2016.03.109

DeLorenzo, M. E., & Fleming, J. (2008). Individual and mixture effects of selected pharmaceuticals and personal care products on the marine phytoplankton species Dunaliella tertiolecta. Archives of Environmental Contamination and Toxicology, 54, 203–210. https://doi.org/10.1007/s00244-007-9032-2

Eea, E. E. A. (1999). Environment in the European Union at the turn of the century. Environmental Assessment Report, 1–446.

Ellesat, K. S., Tollefsen, K. E., Åsberg, A., Thomas, K. V., & Hylland, K. (2010). Cytotoxicity of atorvastatin and simvastatin on primary rainbow trout (Oncorhynchus mykiss) hepatocytes. Toxicology in Vitro, 24, 1610–1618. https://doi.org/10.1016/j.tiv.2010.06.006

Endo, A., Kuroda, M., & Tanzawa, K. (1976a). Competitive inhibition of 3-hydroxy-3-methylglutaryl coenzyme a reductase by ML-236A and ML-236B fungal metabolites, having hypocholesterolemic activity. FEBS Letters, 72, 323–326. https://doi.org/10.1016/0014-5793(76)80996-9

Endo, A., Kuroda, M., & Tsujita, Y. (1976b). ML-236A, ML-236B, and ML-236C, new inhibitors of cholesterogenesis produced by Penicillium citrinium. The Journal of Antibiotics, 29, 1346–1348. https://doi.org/http://dx.doi.org/10.7164/antibiotics.29.1346

Fang, L., & Miller, Y. I. (2012). Emerging applications for zebrafish as a model organism to study oxidative mechanisms and their roles in inflammation and vascular accumulation of oxidized lipids. Free Radical Biology and Medicine, 53, 1411–1420. https://doi.org/10.1016/j.freeradbiomed.2012.08.004

Fent, K., Weston, A. A., & Caminada, D. (2006). Ecotoxicology of human pharmaceuticals. Aquatic Toxicology, 76, 122–159. https://doi.org/10.1016/j.aquatox.2005.09.009

Ferreira, F., santos, M. M., Castro, L. F. C., Reis-Henriques, M. A., Lima, D., Vieira, M. N., Monteiro, N. M., 2009. Vitellogenin gene expression in the intertidal blenny Lipophrys pholis: A new sentinel species for estrogenic chemical pollution monitoring in the European Atlantic coast? Comparative Biochemistry and Physiology C-Toxicology & Pharmacology, 149: 58–64.

Fraher, D., Sanigorski, A., Mellett, N. A., Meikle, P. J., Sinclair, A. J., & Gibert, Y. (2016). Zebrafish Embryonic Lipidomic Analysis Reveals that the Yolk Cell Is Metabolically Active in Processing Lipid. Cell Reports, 14, 1317–1329. https://doi.org/10.1016/j.celrep.2016.01.016

Funatsu, T., Kakuta, H., Tanaka, H., Arai, Y., Suzuki, K., & Miyata, K. (2001). [Atorvastatin (Lipitor): a review of its pharmacological and clinical profile]. Nihon Yakurigaku Zasshi. Folia Pharmacologica Japonica, 117, 65–76. https://doi.org/10.1254/FPJ.117.65

Gee, R. H., Spinks, J. N., Malia, J. M., Johnston, J. D., Plant, N. J., & Plant, K. E. (2015). Inhibition of prenyltransferase activity by statins in both liver and muscle cell lines is not causative of cytotoxicity. Toxicology, 329, 40–48. https://doi.org/10.1016/j.tox.2015.01.005

Gouni-Berthold, I., Berthold, H. K., Gylling, H., Hallikainen, M., Giannakidou, E., Stier, S., Ko, Y., Patel, D., Soutar, A.K., Seedorf, U., Mantzoros, C.S., Plat, J., Krone, W. (2008). Effects of ezetimibe and/or simvastatin on LDL receptor protein expression and on LDL receptor and HMG-CoA reductase gene expression: A randomized trial in healthy men. Atherosclerosis, 198, 198–207. https://doi.org/10.1016/j.atherosclerosis.2007.09.034

Grung, M., Heimstad, M.S., Moe, M., Schlabach, M., Svenson, A., Thomas, K., Woldegiorgis, A., 2007. Human and Veterinary Pharmaceuticals, Narcotics, andPersonal Care Products in the Environment. SFT Report (Oslo), TA-2325/2007,98 pp.

Guo, R., & Chen, J. (2015). Assessing the impacts of dimethoate on rotifers’ reproduction through the pre-exposure history. Ecotoxicology and Environmental Safety, 111, 199–205. https://doi.org/10.1016/j.ecoenv.2014.10.023

Hamadeh, H.K., Bushel, P.R., Jayadev, S., Martin, K., DiSorbo, O., Sieber, S., Bennett, L., Tennant, R., Stoll, R., Barrett, J.C., Blanchard, K., Paules, R.S., & Afshari, C.A. 2002. Gene expression analysis reveals chemical-specific profiles. Toxicological Sciences, 67, 219–231. https://doi.org/10.1093/toxsci/67.2.219

Igel, M., Sudhop, T., & VonBergmann, K. (2001). Metabolism and drug interactions of 3-hydroxy-3-methylglutaryl coenzyme A-reductase inhibitors (statins). European Journal of Clinical Pharmacology, 57, 357–364. https://doi.org/10.1007/s002280100329

Infarmed, I. P. (n.d.). Estatistica do medicamento anteriores – Infarmed – INFARMED, I.P. Retrieved January 31, 2018, from: http://www.infarmed.pt/web/infarmed/infarmed?p_p_id=101&p_p_lifecycle=0&p_p_state=maximized&p_p_mode=view&_101_struts_action=%2Fasset_publisher%2Fview_content&_101_returnToFullPageURL=%2F&_101_assetEntryId=851513&_101_type=content&_101_urlTitle=estatistic

Istvan, E. (2003). Statin inhibition of HMG-CoA reductase: A 3-dimensional view. Atherosclerosis Supplements, 4, 3–8. https://doi.org/10.1016/S1567-5688(03)00003-5

Kasprzyk-Hordern, B., Dinsdale, R. M., & Guwy, A. J. (2009). The removal of pharmaceuticals, personal care products, endocrine disruptors and illicit drugs during wastewater treatment and its impact on the quality of receiving waters. Water Research, 43, 363–380. https://doi.org/10.1016/j.watres.2008.10.047

Key, P. B., Hoguet, J., Reed, L. A., Chung, K. W., & Fulton, M. H. (2008). Effects of the statin antihyperlipidemic agent simvastatin on grass shrimp, Palaemonetes pugio. Environmental Toxicology, 23, 153–160. https://doi.org/10.1002/tox.20318

Kim, B. M., Rhee, J. S., Hwang, U. K., Seo, J. S., Shin, K. H., & Lee, J. S. (2015). Dose- and time-dependent expression of aryl hydrocarbon receptor (AhR) and aryl hydrocarbon receptor nuclear translocator (ARNT) in PCB-, B[a]P-, and TBT-exposed intertidal copepod Tigriopus japonicus. Chemosphere, 120, 398–406. https://doi.org/10.1016/j.chemosphere.2014.07.099

LaLone, C. A., Berninger, J. P., Villeneuve, D. L., & Ankley, G. T. (2014). Leveraging existing data for prioritization of the ecological risks of human and veterinary pharmaceuticals to aquatic organisms. Philosophical Transactions of the Royal Society of London. Series B, Biological Sciences, 369, 20140022-. https://doi.org/10.1098/rstb.2014.0022

Landgraf, K., Schuster, S., Meusel, A., Garten, A., Riemer, T., & Schleinitz, D. (2017). Short-term overfeeding of zebrafish with normal or high-fat diet as a model for the development of metabolically healthy versus unhealthy obesity. BMC Physiology, 17, 1–10. https://doi.org/10.1186/s12899-017-0031-x

Lapworth, D. J., Baran, N., Stuart, M. E., & Ward, R. S. (2012). Emerging organic contaminants in groundwater: A review of sources, fate and occurrence. Environmental Pollution, 163, 287–303. https://doi.org/10.1016/j.envpol.2011.12.034

Lawrence, C. (2007). The husbandry of zebrafish (Danio rerio): A review. Aquaculture, 269, 1–20. https://doi.org/10.1016/j.aquaculture.2007.04.077

Lefcort, H., Freedman, Z., House, S., & Pendleton, M. (2008). Hormetic effects of heavy metals in aquatic snails: Is a little bit of pollution good? EcoHealth, 5, 10–17. https://doi.org/10.1007/s10393-008-0158-0

Livak, K.J., Schmittgen, T.D., 2001. Analysis of Relative Gene Expression Data Using Real-Time Quantitative PCR and the 2-ΔΔCT Method. Methods, 25, 402–408. https://doi.org/10.1006/meth.2001.1262

Lyssimachou, A., Santos, J. G., André, A., Soares, J., & Lima, D. (2015). The Mammalian “Obesogen” Tributyltin Targets Hepatic Triglyceride Accumulation and the Transcriptional Regulation of Lipid Metabolism in the Liver and Brain of Zebrafish. PloS One, 10, 1–2 https://doi.org/10.1371/journal.pone.0143911

Meador, J. P., Sommers, F. C., Cooper, K. A., & Yanagida, G. (2011). Tributyltin and the obesogen metabolic syndrome in a salmonid. Environmental Research, 111, 50–56. https://doi.org/10.1016/j.envres.2010.11.012

Miao, X.-S., & Metcalfe, C. D. (2003). Determination of cholesterol-lowering statin drugs in aqueous samples using liquid chromatography–electrospray ionization tandem mass spectrometry. Journal of Chromatography A, 998, 133–141. https://doi.org/10.1016/S0021-9673(03)00645-9

Moghadasian, M. H. (1999). Clinical pharmacology of 3-hydroxy-3-methylglutaryl coenzyme A reductase inhibitors. Life Sciences, 65, 1329–1337. https://doi.org/10.1016/S0024-3205(99)00199-X

Mu, X., Wang, K., Chai, T., Zhu, L., Yang, Y., Zhang, J., Pang, S., Wang, C., Li, X. (2015). Sex specific response in cholesterol level in zebrafish (Danio rerio) after long-term exposure of difenoconazole. Environmental Pollution, 197, 278–286. https://doi.org/10.1016/j.envpol.2014.11.019

Nash, R. D. M., Valencia, A. H., Geffen, A. J., & Meek, A. (2006). The Origin of Fulton’s Condition Factor — Setting the Record Straight. Fisheries, 31, 236–238. https://doi.org/10.1016/j.anbehav.2015.04.024

Neuparth, T., Martins, C., Santos C. B. de los, Costa, M. H., Martins, I., Costa, P. M., & Santos, M. M. (2014). Hypocholesterolaemic pharmaceutical simvastatin disrupts reproduction and population growth of the amphipod Gammarus locusta at the ng/L range. Aquatic Toxicology, 155, 337–347. https://doi.org/10.1016/j.aquatox.2014.07.009

Nicolás Vázquez, I., Rodríguez-Núñez, J. R., Peña-Caballero, V., Ruvalcaba, R. M., & Aceves-Hernandez, J. M. (2017). Theoretical and experimental study of fenofibrate and simvastatin. Journal of Molecular Structure, 1149, 683–693. https://doi.org/10.1016/j.molstruc.2017.08.044

Oberemm, A. (2000). The use of a refined zebrafish embryo bioassay for the assessment of aquatic toxicity. Lab Animal, 29, 32–40.

OECD. (2013). Test No. 236: Fish Embryo Acute Toxicity (FET) Test. OECD Guidelines for the Testing of Chemicals, Section 2, OECD Publishing, (July), 1–22. https://doi.org/10.1787/9789264203709-en

Ottmar, K. J., Colosi, L. M., & Smith, J. A. (2012). Fate and transport of atorvastatin and simvastatin drugs during conventional wastewater treatment. Chemosphere, 88, 1184–1189. https://doi.org/10.1016/j.chemosphere.2012.03.066

Pereira, A. M. P. T., Silva, L. J. G., Lino, C. M., Meisel, L. M., & Pena, A. (2016). Assessing environmental risk of pharmaceuticals in Portugal: An approach for the selection of the Portuguese monitoring stations in line with Directive 2013/39/EU. Chemosphere, 144, 2507–2515. https://doi.org/10.1016/j.chemosphere.2015.10.100

Pereira, A. M. P. T., Silva, L. J. G., Meisel, L. M., Lino, C. M., & Pena, A. (2015). Environmental impact of pharmaceuticals from Portuguese wastewaters: geographical and seasonal occurrence, removal and risk assessment. Environmental Research, 136, 108–119. https://doi.org/10.1016/j.envres.2014.09.041

Ribeiro, S., Torres, T., Martins, R., & Santos, M. M. (2015). Toxicity screening of diclofenac, propranolol, sertraline and simvastatin using danio rerio and paracentrotus lividus embryo bioassays. Ecotoxicology and Environmental Safety, 114, 67–74. https://doi.org/10.1016/j.ecoenv.2015.01.008

Rodrigues, P., Reis-Henriques, M.A., Campos, J., Santos, M.M., 2006. Urogenital Papilla feminization in male Pomatoschistus minutus from two estuaries in northwestern Iberian Peninsula. Marine Environmental Research, 62, S258–S262. https://doi.org/10.1016/j.marenvres.2006.04.032

Rudling, M., Angelin, B., Ståhle, L., Reihnér, E., Sahlin, S., Olivecrona, H., Björkhem, I., Einarsson, C. (2002). Regulation of hepatic low-density lipoprotein receptor, 3-hydroxy-3-methylglutaryl coenzyme A reductase, and cholesterol 7α-hydroxylase mRNAs in human liver. Journal of Clinical Endocrinology and Metabolism, 87, 4307–4313. https://doi.org/10.1210/jc.2002-012041

Santos, M. M., Ruivo, R., Lopes-Marques, M., Torres, T., de los Santos, C. B., Castro, L. F. C., & Neuparth, T. (2016). Statins: An undesirable class of aquatic contaminants? Aquatic Toxicology, 174, 1–9. https://doi.org/10.1016/j.aquatox.2016.02.001

Sárria, M. P., Soares, J., Vieira, M. N., Santos, M. M., Monteiro, N. M. (2011). Rapid-behaviour responses as a reliable indicator of estrogenic chemical toxicity in zebrafish juveniles. Chemosphere, 85, 1543–1547. https://doi.org/10.1016/j.chemosphere.2011.07.048

Schwartz, D. M., & Wolins, N. E. (2007). A simple and rapid method to assay triacylglycerol in cells and tissues. Journal of Lipid Research, 48, 2514–20. https://doi.org/10.1194/jlr.D700017-JLR200

Sehayek, E., Butbul, E., Avner, R., Levkovitz, H., & Eisenberg, S. (1994). Enhanced cellular metabolism of very low density lipoprotein by simvastatin. A novel mechanism of action of HMG-CoA reductase inhibitors. European Journal of Clinical Investigation 24, 173–178. https://doi.org/10.1111/j.1365-2362.1994.tb00984.x

Soares, J., Coimbra, A. M., Reis-Henriques, M. A., Monteiro, N. M., Vieira, M. N., Oliveira, J. M. A., Guedes-Dias, P., Fontaínhas-Fernandes, A., Parra, S.S., Carvalho, A.P., Castro, L.F.C., Santos, M. M. (2009). Disruption of zebrafish (Danio rerio) embryonic development after full lifecycle parental exposure to low levels of ethinylestradiol. Aquatic Toxicology, 95, 330–338. https://doi.org/10.1016/j.aquatox.2009.07.021

Sousa, M. A. D. de. (2013). Analysis of pharmaceutical residues in wastewaters, surface and drinking waters – study of the removal efficiency through conventional and advanced treatment processes. Faculdade de Farmácia da Universidade do Porto.

Stebbing, A. (1982). Hormesis – the Stimulation of Growth By Low-Levels of Inhibitors. Science of the Total Environment, 22, 213–234. https://doi.org/10.1016/0048-9697(82)90066-3

Thorpe, J. L., Doitsidou, M., Ho, S., Raz, E., & Farber, S. A. (2004). Germ Cell Migration in Zebrafish Is Dependent on HMGCoA Reductase Activity and Prenylation Short Article. Developmental Cell, 6, 295–302. https://doi.org/10.1016/S1534-5807(04)00032-2

Torres, T., Cunha, I., Martins, R., & Santos, M. M. (2016). Screening the toxicity of selected personal care products using embryo bioassays: 4-MBC, propylparaben and triclocarban. International journal of Molecular Sciences, 17, 1762. https://doi.org/10.3390/ijms17101762

van der Wulp, M. Y. M., Verkade, H. J., & Groen, A. K. (2013). Regulation of cholesterol homeostasis. Molecular and Cellular Endocrinology, 368, 1–16. https://doi.org/10.1016/j.mce.2012.06.007

Vandenberg, L. N., Colborn, T., Hayes, T. B., Heindel, J. J., Jacobs, D. R., Lee, D.-H., Shioda, T., Soto, A.M., vom Saal, F.S., Zoeller, W.V.W.R.T., Myers, J. P. (2012). Hormones and endocrine-disrupting chemicals: low-dose effects and nonmonotonic dose responses. Endocrine Reviews, 33, 378–455. https://doi.org/10.1210/er.2011-1050

Vergauwen, L., Benoot, D., Blust, R., & Knapen, D. (2010). Long-term warm or cold acclimation elicits a specific transcriptional response and affects energy metabolism in zebrafish. Comparative Biochemistry and Physiology, Part A, 157, 149–157. https://doi.org/10.1016/j.cbpa.2010.06.160

Verlicchi, P., Al Aukidy, M., & Zambello, E. (2012). Occurrence of pharmaceutical compounds in urban wastewater: Removal, mass load and environmental risk after a secondary treatment-A review. Science of the Total Environment, 429, 123–155. https://doi.org/10.1016/j.scitotenv.2012.04.028

